# What can the current global bacterial taxonomy knowledge reveal about the actual (past and present) ecological success of each lineage?

**DOI:** 10.1101/2025.09.20.677513

**Authors:** Andrea Squartini

## Abstract

Our up-to-date sources of the man-known microbiology are publicly available, and include, as the most complete repository, the Genome Taxonomy Database GTDB https://gtdb.ecogenomic.org/, encompassing, on one end, 196 phyla, across which, at the other end, 107,235 bacterial species have been currently defined. The intermediate ranks feature 541 classes, 1,863 orders, 4,896 families, and 23,112 genera. Such a dataset is hereby used, not as a source of DNA sequences but rather as the handiest and presently most exhaustive checklist matrix of our current taxonomical notions.

In the mathematical ‘zero assumption’ of an equal opportunity chance for each taxon throughout a ‘parallel’ evolution, a ‘perfectly even’ community featuring those values would display today, within each of the 196 phyla, the mean values that would result by dividing the actual observed ones by the number of phyla. Such a virtual assemblage would therefore have the following numbers per phylum in each of them: 2.76 classes, 9.51 orders, 24.98 families, 117.92 genera, and 547.12 species. The deltas between these set-point null hypothesis values and the truly observed and described numbers for each lineage member featured by the database (numbers of species per genus, genera per family, families per order, orders per class, and classes per phylum) can reveal how much, and in which lineages, their evolution differed from each of those mere ranks’ means. The comparison among the actual data also allows one to answer a series of questions: 1) What is the actually occurred uneven dynamics of evolutionary differentiation across each lineage?; 2) Which taxa are, at present, the ‘winners’ of the evolutionary race?, And, have they always led the run, or when did they overtake their formerly faster competitors? 3) Are there chronological bottlenecks that violate the assumption of either a constant or of a progressive speed of evolution (i.e. is speciation to each deeper, and more ‘modern’ rank, consistently faster than its former step, or is there evidence of evolutionary congestion ‘jams’?); 4) which past step turns out to have been the most determinant to shape the modern assemblages? Which ones, among the taxa dominating today’s diversity, have risen up just recently (e.g. by a massive radiation from genera), and which ones had instead started or accomplished the critical expansion that still keeps them as leaders, in remote times (e.g. by class or order radiations?; 5) Which patterns characterize in this respect the lineages of the most relevant bacterial agents in food, soil, industrial, clinical and environmental microbiology?

Corresponding queries are addressed for the 5,869 species of Archaea.

The methods and the results of this database-mined in silico analysis are hereby presented and discussed. The dataset provided as supplementary material allows the reader to address the above aspects for any taxon of interest within the 107,235 species covered by this analysis.

## Introduction

The classification of bacterial life has long been constrained by the limitations of phenotype-based taxonomy and the partial resolution offered by 16S rRNA gene sequencing. These approaches laid the groundwork for microbial taxonomy and for decades have been its gold standards. However, although foundational, they often failed to reflect true evolutionary relationships, resulting in polyphyletic groupings and inconsistent taxonomic ranks (Yarza et al., 2014; Beiko, 2015). In response to these and other challenges, Parks et al. (2018) proposed a novel phylogeny, which was inferred upon concatenating 120 ubiquitous proteins, generating a database that by now has grown to encompass 732,475 genomes. The Taxonomic groups in such a curated classification describe monophyletic lineages with similar phylogenetic depth after having normalized the data for lineage-specific rates of evolution.

This introduced a transformative genome-based taxonomy: the Genome Taxonomy Database (GTDB) (Parks et al., 2018), available for downloads at https://gtdb.ecogenomic.org/, which leverages concatenated protein phylogenies and relative evolutionary divergence (RED) to standardize bacterial classification across all available genomic data. This framework not only resolved widespread polyphyly, notably within genera such as *Clostridium* and *Bacillus*, but also redefined higher-order taxa, including the subdivision of Proteobacteria and the amalgamation of the Candidate Phyla Radiation into the single phylum Patescibacteria. Since its inception, the GTDB taxonomy has catalyzed a paradigm shift in microbial systematics. Subsequent studies have expanded and refined its principles. For example, Barco et al. (2020) proposed a genus demarcation strategy based on average nucleotide identity (ANI) and genome alignment fraction, offering a robust complement to GTDB’s RED-based rank normalization.

Similarly, Lalucat and coworkers (2020) applied phylogenomic methods to reassess the genus *Pseudomonas*, demonstrating how whole-genome analyses can reconcile taxonomic inconsistencies and uncover cryptic diversity within well-studied lineages. These developments underscore a growing consensus: genome-based metrics provide a more coherent and evolutionarily meaningful framework for bacterial taxonomy than traditional marker gene approaches.

The impact of GTDB has been profound and far-reaching. Subsequent tools from the same group like GTDB-Tk (Chaumeil et al., 2019) have improved the computing capability of the system and enabled automated and scalable taxonomic classification of bacterial and archaeal genomes, facilitating the integration of metagenome-assembled genomes (MAGs) into the taxonomic framework. This is essential for environmental microbiology, where uncultured organisms dominate and traditional classification methods fall short.

Subsequently Liao et al (2020) focused on refining or reclassifying orders in particular bacterial groups, correcting misplacements under older taxonomies. They used the Parks et al. (2018) tree as a reference for what should be considered monophyletic, and to disentangle related issues.

Improvements of the system have been produced (Parks et al.,2022), where updated versions of the Genome Taxonomy Database were provided, amply expanding genome sampling (bacteria + archaea), improving species clusters, improving tools for tracking taxonomic changes, and genome quality. These were directly built from the first paper framework, adopting the same phylogenomic-based tree, the same rank normalization by relative evolutionary divergence, including many more genomes, better metadata, and tooling. Later, the group added 329 higher-rank taxa that were previously unnamed (or bore a placeholder), helping to solidify classification and nomenclature under the GTDB logic (Chuvochina et al., 2023). This addition directly leveraged rank-normalized taxonomy, using its definitions for higher ranks, monophyly.

Since the publication of the original GTDB study, several subsequent works have built upon its framework. These follow-ups have applied its methods to broader datasets, tested its proposed revisions in other bacterial groups, or used its findings as a reference point for comparative taxonomy in different environments. For example, the MiDAS 5 project (Dueholm et al., 2024) widely expanded full-length 16S rRNA gene sequence databases, enabling improved species- and genus-level classification in complex microbial communities (e.g., anaerobic digesters), and highlighted how much undiscovered diversity remains even in relatively well-studied environments, validating the need for more genome-scale taxonomic approaches.

Other pieces of research (Prinzi et al., 2023) have reviewed evolving rules and challenges in bacterial nomenclature and taxonomic practice, underscoring that, even as genomic data accumulates, issues of standardization, reproducibility, and nomenclatural stability remain unresolved. These developments collectively suggest that the field is heading toward a more integrative taxonomy, one that combines genomic, phenotypic, ecological, and evolutionary data, but also that key gaps remain, both in method and in consensus.

The GTDB genome-based taxonomy represented both a methodological advance, employing large-scale phylogenomics, well-defined rank normalization, and better handling of uncultured diversity— and a conceptual shift: the recognition that taxonomies should increasingly reflect evolutionary history as inferred from genomes, not just a patchwork of phenotypic traits and a small number of marker genes. Since 2018, several lines of work have further validated, refined, or applied this framework, as well as raised new challenges.

- Refinements in taxonomic resolution and application to specific lineages: for example, Li et al. (2024) undertook “Phylogenomic analyses and reclassification of the *Mesorhizobium* complex,” proposing nine novel genera and reclassifying fifteen species, using the kinds of overall genome-relatedness indices that are consistent with the GTDB‐style approach. This demonstrates how genome-scale approaches allow resolving fine-scale relationships that older taxonomies may have conflated or mis-assigned.
- Increased requirement of genomic metrics and standards: beyond just phylogeny, newer studies are more frequently using average nucleotide identity (ANI), digital DNA–DNA hybridization (dDDH), and other genomic similarity metrics as standard corroborative tools for distinguishing species or genera. Those are tools whose importance was suggested but less uniformly applied in 2018. Also, the minimal standards for genome-based species descriptions have been more precisely articulated and increasingly enforced in many journals (Sant’Anna et al., 2020).
- Practical and nomenclatural challenges: As taxonomy changes, there are consequences: databases, diagnostic labs, epidemiology, and clinical microbiology are all affected when familiar taxa are renamed or re-grouped. This has led to work summarizing taxonomic changes (for example, in animal and domestic species from 2018–2021, documenting novel taxa and revised status) to monitor how taxonomy is evolving in practice (Munson et al., 2023).

Another report focused on how taxonomic proposals/reclassifications (especially genome-based ones) should be validated in light of new data, standardization, reproducibility etc., which raises issues about how consistent and stable genome-based taxonomy frameworks are, under various biases, as sampling, assembly quality, etc., emphasizing areas where further rigor is needed, standards for naming, rules for handling uncultured organisms, consideration of phenotypic/ecological data in addition to genome data (Venter et al, 2024).

Another study (Shoguchi et al., 2024) applied the GTDB-Tk toolkit to classify genomes from a marine environment, then used those classifications to draw ecological or functional conclusions. This demonstrates the practical downstream utility of the genomic taxonomy. It constituted an advancement of the applied side: i.e., not just proposing taxonomy, but using it for interpreting ecological or metabolic functional data.

Recent updates in prokaryotic taxonomy include a thorough report for the archaea in which the authors elaborated an improved annotation using the genome taxonomy database and RNAseq data (Grant et al., 2025). It is also important to mention, as a companion metadata repository, the updated Bac Dive database (Schober et al., 2025).

Notwithstanding the progress of our knowledge in microbial systematics, important issues persist: how to best normalize taxonomic rank boundaries (how much evolutionary divergence is “enough” to call something a phylum vs class, etc.); how to ensure stability versus frequent revisions (which can make data comparisons over time difficult); how to deal with taxa represented only by uncultured genomes; how to integrate phenotype, ecology, and genome in a balanced taxonomy. Also, the computational and sampling biases (which environments are sequenced, and which genomes are high quality) can skew our understanding of diversity and phylogeny.

A further necessary premise is that our knowledge in microbial taxonomy, unlike the cases of zoology and botany, cannot count on a consistent availability of fossils, and is essentially grounded on molecular data, which are assumed to reflect extant taxa. For what regards extinction, the estimation of the proportions of lost species, therefore, relies only on theoretical modeling. Nevertheless, the extrapolations concur on a scenario in which extinction rates for microbes would practically be negligible, due to the ubiquity, dispersiveness, resistance, and resilience of their cells (Staley, 1997; Weinbauer and Rassoulzadegan, 2003). Although hypotheses are debated (Louca, et al., 2018), so far the current information allows to consider our acquired knowledge on microbial phylogeny as not being particularly affected by the disappearance of lineages. This allows one to speculate on the actual evolutionary success or stalling of all groups as inferable from their efficiency of radiation within the tree of life, which is the object of the present analysis. Nevertheless, the possible extinction background, whatever its burden would be, constitutes a part of the multiple factors (some with a positive and some with a negative sign), that contribute just as well to the results that are measured in this data elaboration, i.e. to the taxa numbers in each rank and to their reciprocal proportions.

In the present work, the GTDB repository was used solely as a source of the known taxonomical names and their corresponding lineages. The idea applied here was the following: a bacterial taxonomy table in a spreadsheet, once its last column (species) is cleaned from redundancies by eliminating duplicate names and leaving only unique ones, contains the full array of current bacterial knowledge arranged in six distinct columns (phylum, class, order, family, genus, species). At that point, the subsequent elaboration, leading to all ranks’ numbers and their proportions, is just a matter of counting them with suitable formulas. Differences in such proportions across groups can highlight hitherto unknown differences in the phylogenetic dynamics of each lineage.

## Methods

The bacterial and archaeal taxonomy datasets (Release v220.0; bac120_taxonomy.tsv and ar53_taxonomy.tsv) were downloaded from GTDB at https://gtdb.ecogenomic.org/. All subsequent operations were carried out in Microsoft Excel. The starting tables (in tsv format) contained the lineages for each accession in rows whose columns feature the text values of names for the following ranks: domain, phylum, class, order, family, genus, species. The database used for bacteria featured 584,382 entries in as many rows, which encompassed all the accessions (including those of different strains bearing the same species names). In order to obtain a dataset with unique species names and quantify all their respective higher ranks, the following elaborations were applied in the spreadsheet. Extra columns for counting values were introduced on the right of each original column (e.g., =COUNTIF($B$1:$B$584382;B584382). Unique values were counted for each text-containing column by the formula, e.g., =ROWS(UNIQUE(A1:A584382). Such a row yielded the numbers of unique known names for phyla, classes, orders, families, genera, and species. Next, duplicate species names (and connected rows with their lineage ranks text values across the worksheet) were eliminated by selecting the species column and using the command Data>Remove duplicates. The resulting datasets featured the 107,235 rows corresponding, non-redundantly, to the known named species for bacteria, and the 5,869 ones for archaea. Subsequently, the strategy to obtain the numbers of genera per family, those of families per order, orders per class, and classes per phylum was to progressively remove the rightmost rank columns (e.g. first the two columns containing names and counts of the species, thus obtaining a worksheet in which the remaining rows had in that case only all genera, and in which, the above-mentioned COUNTIF formula automatically updated its counts to the remaining columns’ values. Saving the new worksheet with a different name and using it. in turn. as a source to repeat this operation, working each time leftwards, always for the two rightmost columns, allowed to obtain the numbers for all the other ranks in five separate resulting worksheets. All the respective columns of interest (number of classes per phylum, numbers of orders per class, etc.) were then copied, and their values were pasted into a new final and collective worksheet in which the resulting data can be inspected (Supplementary Dataset S1 Bacterial and Archaeal elaborated counts.xslx).

## Results and Discussion

### Overall rank proportions

The results of all the elaborations done in the present analysis are compiled in the Microsoft Excel file of the supplementary dataset. It contains two worksheets (bacteria and archaea). The taxa in each set of columns are ordered by the number of species in decreasing abundance from the top. The % column next to each of the species counts columns expresses the percentage of the species listed in each row over the total species number. Aside from the counts of the ranks in each set, there are groups of columns that report the number of members of each further rank, per member of the prior rank (e.g., number of classes per phylum, number of orders per class, etc.).

The global scores for all ranks of bacteria and archaea are shown in the following table

**Tab 1.** Total taxa per rank. number of different unique names featured in each of the six taxonomy ranks in the two domains of Bacteria and Archaea

|  | Bacteria | Archaea |
| --- | --- | --- |
| Phyla | 196 | 21 |
| Classes | 541 | 66 |
| Orders | 1,863 | 167 |
| Families | 4,896 | 564 |
| Genera | 23,112 | 1,847 |
| Species | 107,235 | 5,869 |

To better illustrate how to inspect the datasets with the results of the present elaboration, excerpts of the top rows are clipped and provided in the form of figures (Figs. 1-4) of this report.

**Fig. 1.**
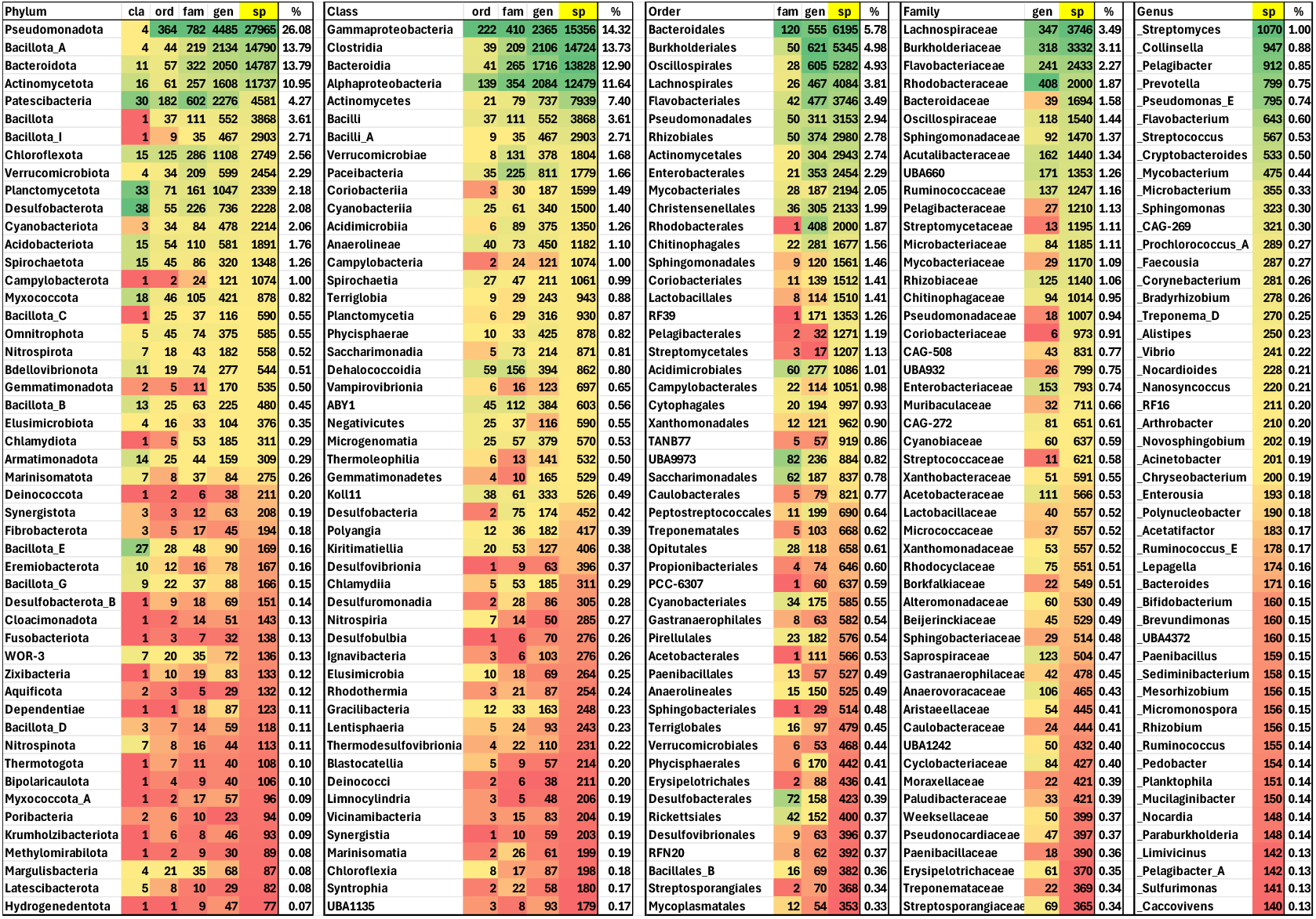
Bacterial dataset. Numerical values. Data report the first 50 rows of the dataset (Supplementary material S1) showing the numbers of the top phyla, classes, orders, families, genera, and the corresponding species number within each category. The values in each rank set are ordered in decreasing abundance by those in their species columns. The derived ratios are presented in the companion Fig.2. The conditional formatting by background color shades (from green to red for decreasing numerical values) is applied by column.

In Fig. 1, from the bacteria worksheet, the top 50 entries of each taxonomy rank are listed. In these tabled images, entries are ordered in decreasing abundance, ranked by the number of species, and the percent value over the total number of species is shown aside.

In Fig.2, in which the excerpt of the corresponding side columns of the Excel dataset (cla/phy, ord/cla, etc..) is pasted, the resulting ratios allow one to assess how evolutionarily productive a rank has been (cla/phy: number of classes per phylum, ord/cla: number of orders per class, etc.). The view enables to compare at which stage of the evolution each group has been mostly prolific. E.g. Pseudomonadota (formerly Proteobacteria) is the phylum with most species, but it is a ‘late branching’ taxon, as it had sprouted only four classes, while Planctomycetota is a strong ‘early branching’ phylum, which has given rise to 33 classes, which however were not equally productive for the stemming of orders, families, and genera, and ended up accounting only 2.181 % of the total bacterial species.

**Fig. 2.**
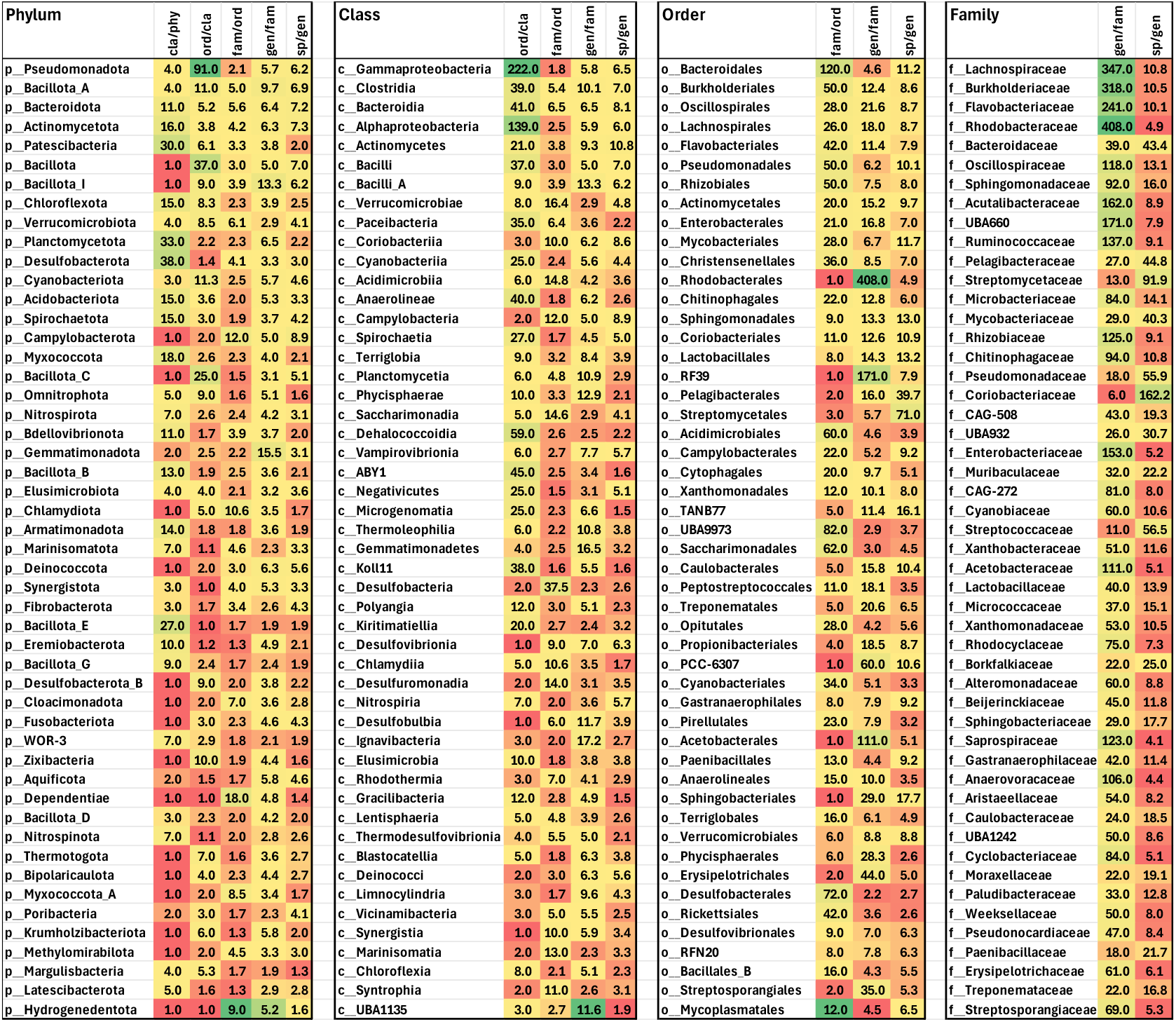
Bacterial dataset; Ratios. Data are derived from those shown in the columns of Fig.1 and show the quotients obtained by dividing each of the values by those of their preceding rank, obtaining the numbers of: classes per phylum, orders per class, families per order, and genera per family (the number of species per genus was already shown in the last column on the right of Fig. 1). The order by which each entry is listed in each set is the same of Fig. 1 being determined by the number of species in each rank group.

As commented in the introduction The GTDB Database, (which does not contain any of these numerical values and was here used simply to acquire the current taxa names and their lineages), features the updated version of the bacterial and archaeal taxonomy which had been revised based on all the markers adopted for the faithful Relative Evolutionary Divergence introduced by Parks et al (2018), in which former terms, as just recalled, have acquired new names and formerly divided groups have been unified (for example, the classes of Proteobacteria are now only 4 (Alpha-, Gamma-, Zeta- and Magnetococcia) with cases as, e.g., the former name Betaproteobacteria having disappeared and its members now being included within Gammaproteobacteria, while members of Deltaproteobacteria were split in different phyla as and Epsilonproteobacteria became the new phylum Campylobacterota, etc., while in the Archaea, the former phylum Crenarchaeota has now become Thermooproteota, etc.).

The same information is provided from the archaea worksheet of the compiled dataset. In Fig. 3 and Fig. 4, the corresponding results, in this case for the top 21 entities (including all archaeal phyla) are shown with the same criteria used for the bacteria.

**Fig. 3.**
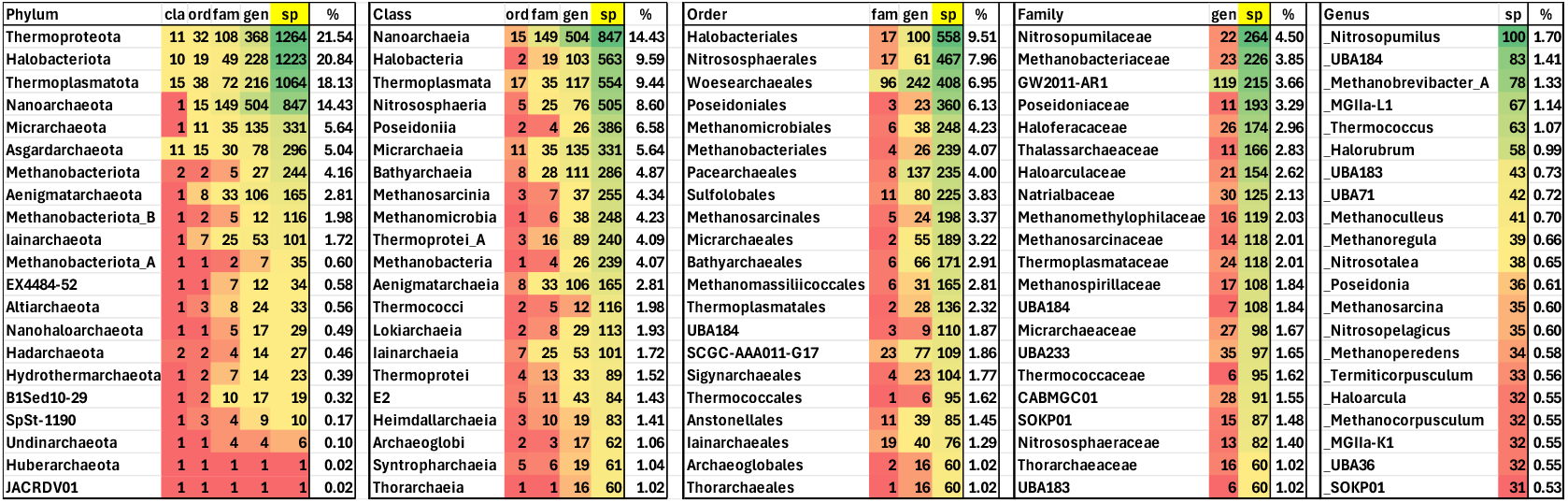
Archaeal dataset. Numerical values. Data report the first 21 rows (covering, for the first rank, all phyla) of the dataset (Supplementary material) showing the numbers of the phyla, and those of the top classes, orders, families, genera, and the corresponding species number within each category. The values in each rank set are ordered in decreasing abundance by those in their species columns. The derived ratios are presented in the companion Fig. 4.

**Fig. 4.**
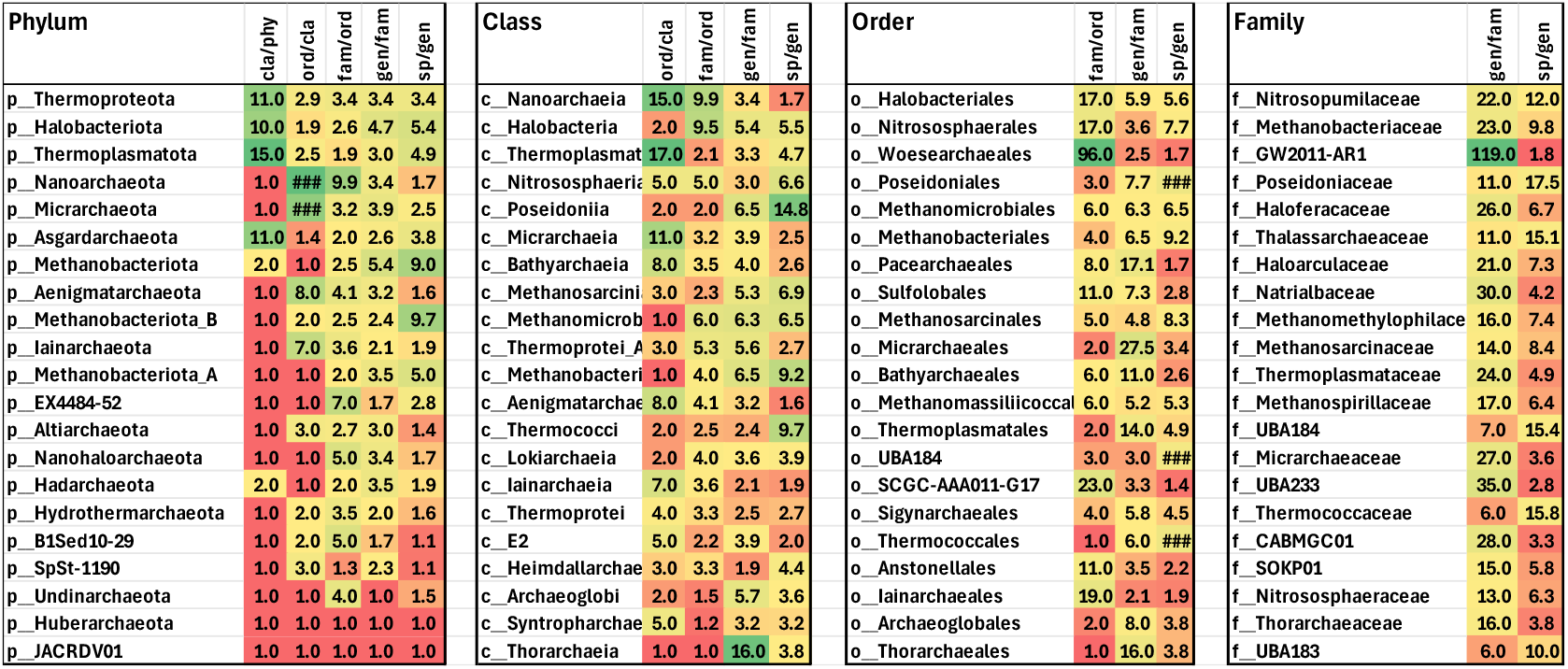
Archaeal dataset; Ratios. Data are derived from those shown in the columns of Fig.3 and display the quotients obtained by dividing each of the values by those of their preceding rank, obtaining the numbers of: classes per phylum, orders per class, families per order, and genera per family (the number of species per genus was already shown in the last column on the right of Fig. 1). The order by which each entry is listed in each set is the same of Fig. 3 being determined by the number of species in each rank group.

### Rank ‘sprouting’ efficiency is step-dependent

Bacteria are currently divided into 196 phyla and 107,235 species. Between these ranks, there are four other ones (class, order, family, genus), through which evolution has unfolded. But, posing a question about its balance, did the five steps occur with equal kinetics? If one were to set a purely theoretical baseline to be used for comparisons with the really observed numbers, in the case of a ‘perfectly additive’ phylogeny, the numerical gap between the number of phyla (the ‘left end’ of the lineage line), and he number of species (the ‘right end’) would present identical mathematical increments at each step. equal to 107,235 - 196 = 107,039, which, divided by five, is 21,407.8.

**<u>196</u> (phyla)** + 21,407.8 = <u>21,603.8</u> classes; + 21,407.8 = <u>43,011.6</u> orders; + 21,407.8 = <u>64,419.4</u>families; + 21,407.8 = 85,827.2 genera; + 21,407.8 = **<u>107</u>**,**<u>235</u> (species)**

If the mathematical function were instead that of a ‘perfectly branching’ radiation process (based on equal multiplications at each step instead of equal additions, in order to reach 107,235 species, from 196 phyla in each of the five ‘jumps’, it would be necessary to multiply each time by 3.528711, which yields the following progression:

**<u>196</u> (phyla)** x 3.528711 = <u>691.6</u> classes; x 3.528711 = <u>2,440.5</u> orders; x 3.528711 = <u>8,612</u> families;x 3.528711 = <u>30,389.3</u> genera; x 3.528711 = **<u>107</u>**,**<u>235</u> (species)**

Such a fixed-coefficient model, which is more coherent with the expected behaviour of biological replicators, yields numbers of ranks much closer to the ones that are observed in the databases, i.e.; 541 classes, 1,863 orders, 4,896 families, 23,112 genera. The level of overestimation with respect to the actually observed data has the following coefficients: 1.28 for classes, 1.31 for orders, 1.76 for families, and 1.31 for genera. For archaea, the corresponding multiplier coefficient value to reach 5,869 species from 21 phyla would be 3.0851

The above considerations allow one to focus more closely on the goals of the present analysis, which aims at measuring the different pace by which each rank underwent a proliferation in the number of its members, both in the overall domains and differentially in each single lineage. The opposite mechanisms that underpin phenomena as biological evolution, where complexity and richness increase with time, are depicted in Fig. 5

**Fig. 5.**
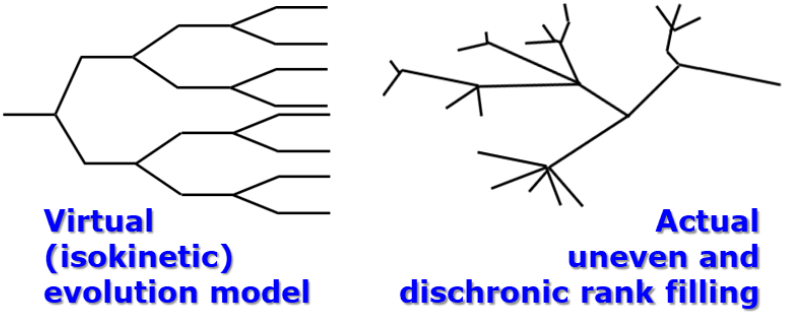
Sketched theoretical vs. realistic types of complexity progression. The left graph represents an environment-independent mathematical function, while the irregular tree on the right assumes the shaping of branches linked to success and failures resulting from genotype by environment interactions.

A tabulated and plotted example of ideal vs. measured meta-taxonomy differences for both bacteria and archaea is provided in the next two figures. The ‘virtual’ numbers that pertain to the imaginary perfectly additive progression of the first example above, to fill the numbers of the four ranks between phyla and species, for the bacteria, are shown in Fig. 6. The numbers panel table compares that theoretical increments to the actual numbers of unique names for each rank that were extracted from the downloaded dataset, which expectedly show a non-linear kinetics.

**Fig. 6.**
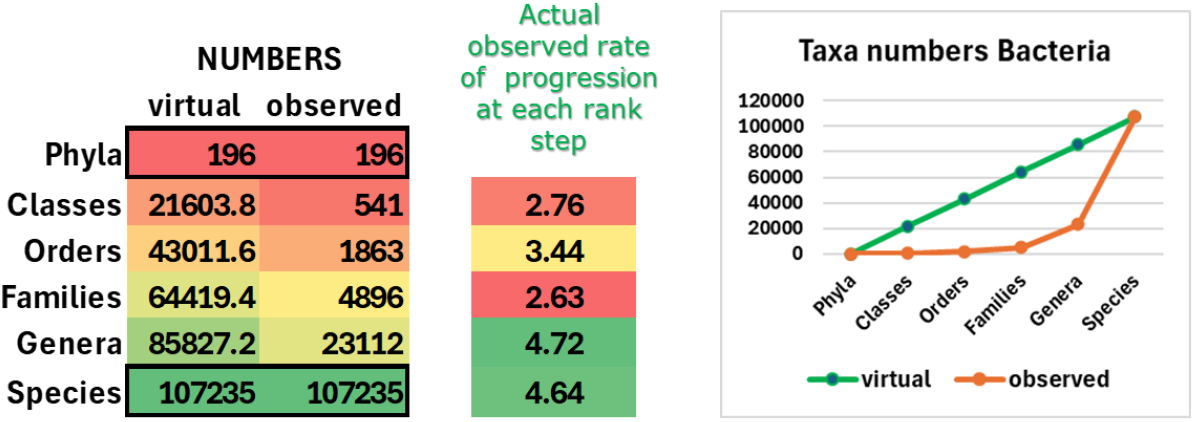
Numbers by ranks and fold progression ratios in bacteria. In the numbers panel, the ‘virtual’ column shows the purely mathematical linear progression to fill the interval between the two extremity ranks (phyla to species) by equal increments. The observed numbers (second column) are the ones extracted from the database in the present work. The third column values (rate of progression) are obtained from the actual observed data of the second column by dividing each value by its preceding one (n. of classes divided by n. of phyla, n. of orders divided by n. of classes, etc. The right panel plots in the line graph the numbers shown in the left table

In the third column, the fold-progression rate column reports the output ratios resulting in each case from the observed numbers of the second column (n. of classes per phylum, n. of orders per class, etc.). Such ratios enable inspection of the efficiency of ‘next rank sprouting’ through the different steps. Fig. 7 reports the same for the archaea.

**Fig. 7.**
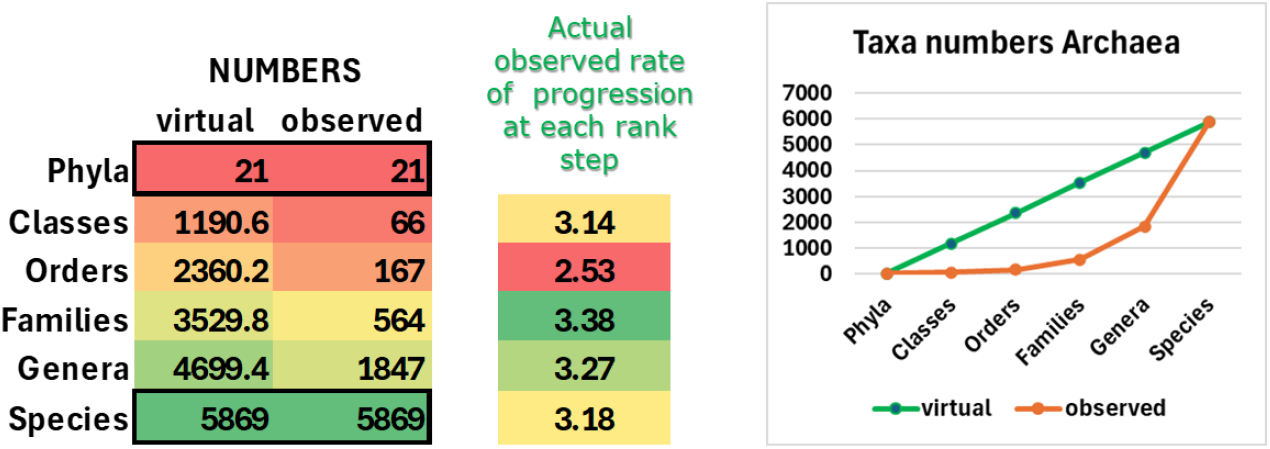
numbers by ranks and fold progression ratios in archaea. The panels follow the same scheme and meaning of those described for bacteria in Fig. 6.

It can be seen that in bacteria, looking at the numbers stemming from the real observed data, the ‘birth rate’ of classes from phyla (2.76) results less intense than that of genera from families (4.72), which in turn is similar to that of species from genera (4.64). The line graphs show the different shape in taxa accumulation between the virtual mathematical linearity and the growth curve that the observed true data define.

In archaea (Fig. 7), on the contrary, the rate of classes per phylum (3.14) was very similar to that of species per genera (3.18).

In archaea (Fig. 7), the same calculations can be made and contrary to the bacterial situation, in the progression quotients column, the rate of classes per phylum (3.14) was very similar to that of species per genus (3.18).

The data in both right panel line graphs show for bacteria and archaea a comparable shape of progression. Which would instead appear divergent when the same data are shown on the same scale, as shown in Fig. 8.

**Fig. 8.**
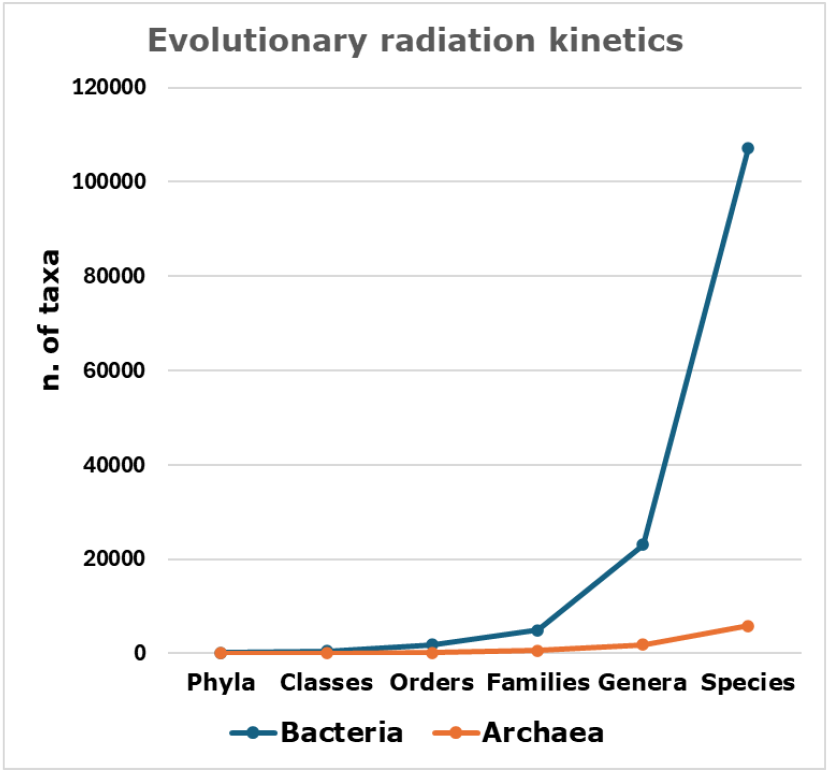
Comparison of the bacterial and archaeal rank richness. The numerical data (true observed numbers) are the same as those shown in Fig. 6 and Fig. 8, but are hereby plotted on the same scale.

The differences are due to the numbers’ disproportions since bacteria differentiate from higher starting numbers, which confer them an inherent mathematical advantage in their progression, not necessarily requiring faster speciation dynamics. Similar to PCR or cells growth in a microbial culture, the same kinetics of doublings, with the same generation time, producing the exponential phase, entails differences of throughput linked, and due, to the difference in the starting numbers. Under this light, the concept of archaea as less prone to evolve would not be particularly supported when judging from the numbers of generated phyla, classes, orders, families, genera and species, as the numbers and the progression are also consistent with being at an earlier point of the exponential curve.

### Evolutionary ‘jams’ and rate plateaus

Using from now on only the real observed data, and continuing to use the ratios of the number of names stemming per member of their higher rank, the plotted graphs show a number of peculiarities. For bacteria and archaea, the resulting shapes are the ones shown in Fig. 9. In these plots, instead of the numbers of members per rank, the quotients of those numbers with respect to the originating level are used, allowing to visualize whether the fold-progression rate towards higher absolute numbers is unidirectionally increasing or shows instead any slowdown at given steps.

**Fig. 9.**
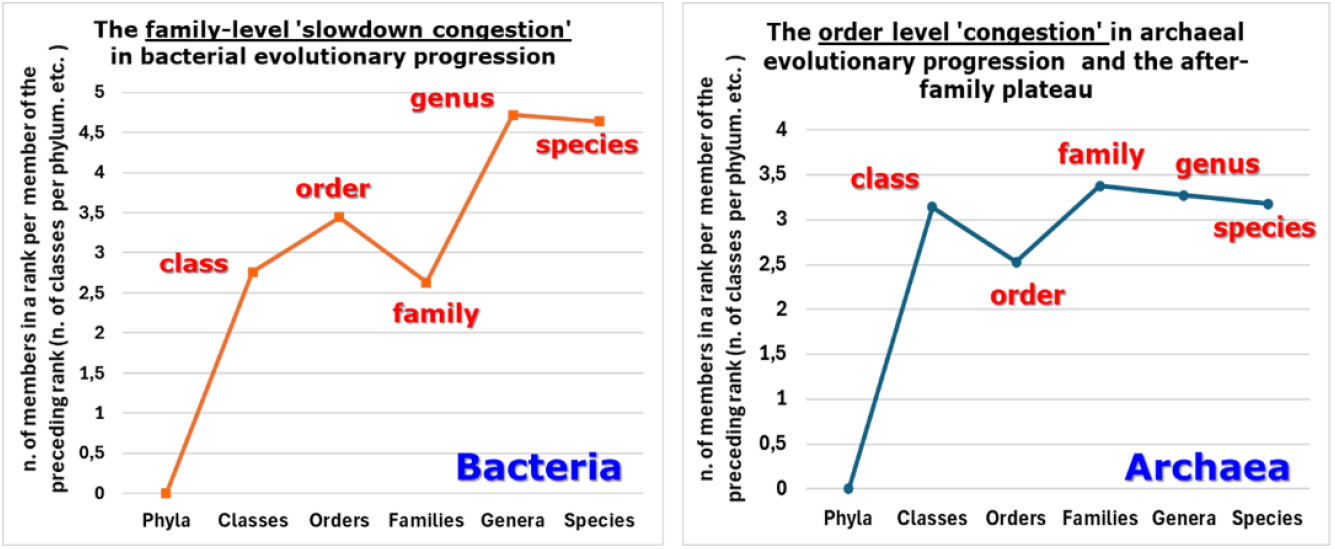
Rank-filling efficiency throughout each transition level. Comparison between bacterial and archaeal evolutionary kinetics in terms of next-rank radiation rate across the six ranks.

As shown in Fig. 9 (left panel), in bacteria, using the all-taxa collective data, the ‘budding rate’ of next-level ranks features a peak at the order level, followed by a ‘bottleneck’ for the family rank and a reprise to the genus. In archaea (Fig. 9, right panel), such a phenomenon results in an anticipation showing its tip at the class level and its minimum at the order rank. In both domains, a slow rate decrease generating a plateau phase is observed, but it starts at the genus level in bacteria, while it is already noticeable from the family level in archaea.

Plotting the two curves in the same graph (Fig. 10) allows to better appreciate these differences.

**Fig. 10.**
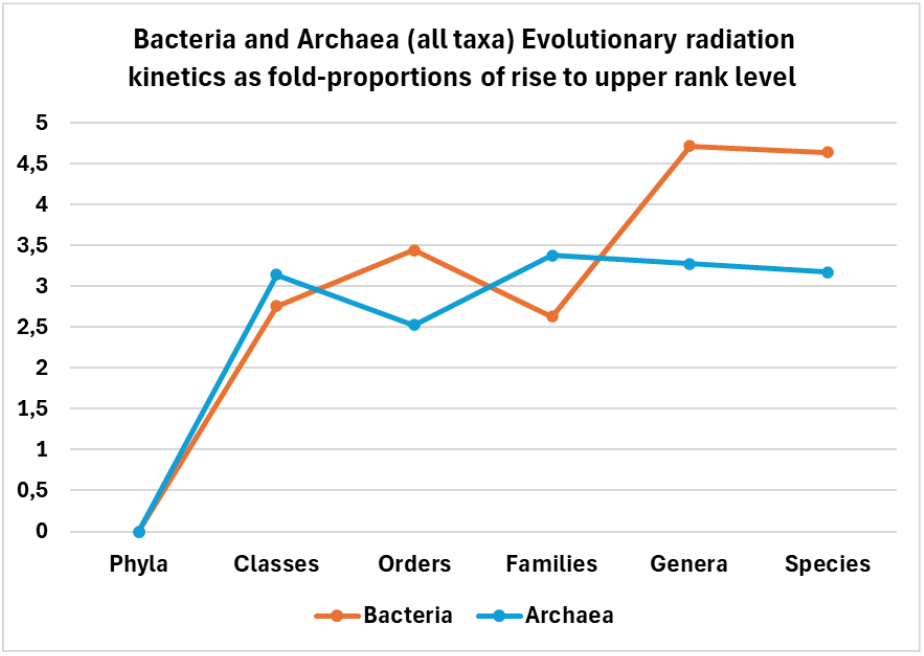
Direct comparison of the rates in the two domains. The curves of bacterial and archaeal taxa accumulation rates shown in Fig. 9 are plotted together to evidence the anticipated slowdown and plateau of the latter group.

### Downwards-upwards alternate fluctuations are due to the ecologically successful (=species-rich) lineages

In the prior paragraph, evidence of rate slowdown was shown to occur characteristically in both bacteria and archaea, and in the latter, the phenomenon was anticipated by one rank. However, those results are the ones that come from the average contribution of all taxa. But the taxonomy structure shows (Fig.1) that in bacteria, the numerically dominant groups in terms of number of encompassed species are relatively very few: just 15 phyla out of 191 account for a percentage > 1% in terms of species, and include collectively 90.39 % of the total species. Another 27 phyla have percentages between 0.1% and 1%, while the real majority (154 phyla) score percentages below 0.1%.

In the archaea domain, the structure is less oligarchic, but, here again, 10 phyla out of a total of 21 have more than 1% of species and account for 96.3 % of the species total. With these differences in mind, when looking separately at the phyla leading the species number scores (15 phyla in bacteria and 10 in archaea), one finds (Fig. 11) that those were the ones in which the phylum-to-class radiation has been the dominant shift, and it is also clearly proportional to the number of species that a phylum encompasses.

**Fig. 11.**
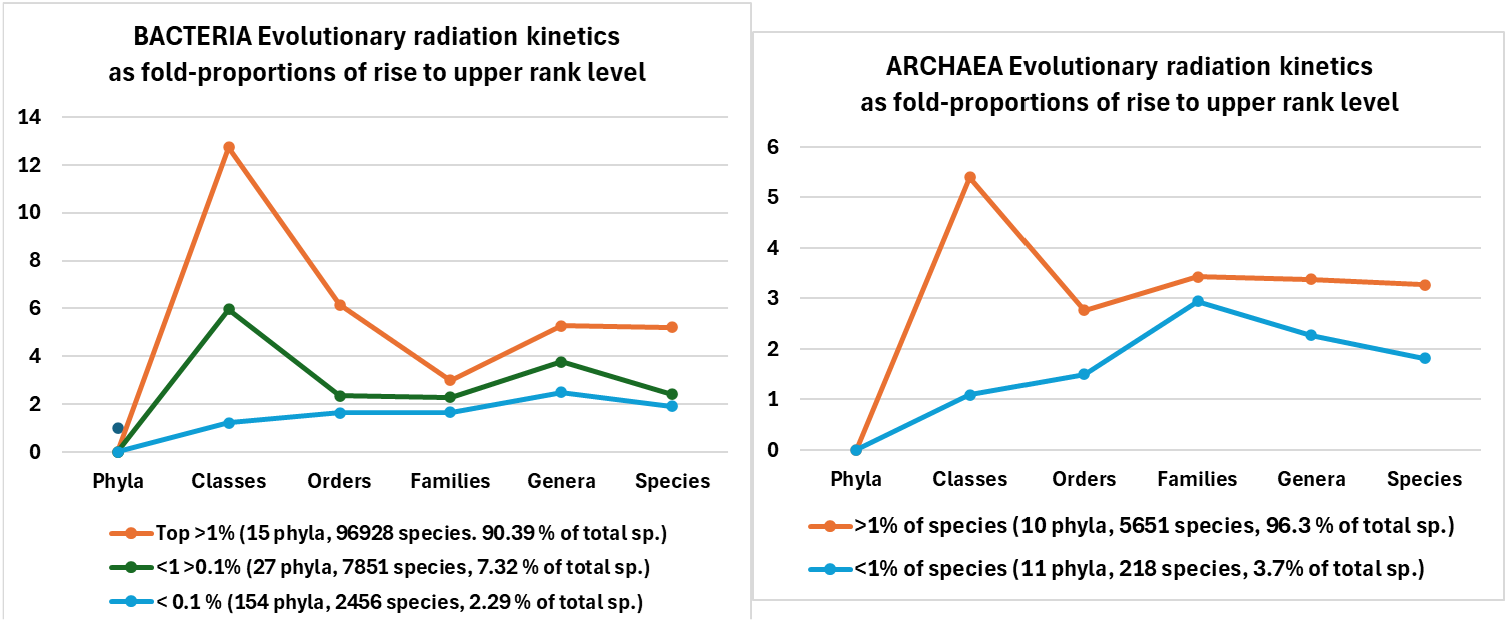
Differential contribution to the radiation rate as a function of species richness. The data used in the graphs shown in Figs. 9 and 10 are here analyzed by splitting the bacterial data into three categories and the archaeal ones into two categories, to reveal the uneven behavior of evolutionarily successful taxa with respect to the unproductive ones.

Therefore, in bacteria (Fig. 11, left panel), the observed tipping point at the order level is actually contributed by the extremely vast proportion of poorly-prolific phyla (154 over a total of 196) that account for only 2.29 % of the total species. In archaea, the corresponding comparison reveals that the order-level decline is made by the species-rich phyla, while the species-poor ones have a peak at family rank and none at class level.

### The rate fluctuation dynamics in rank proliferation are phylum-specific

Using the same metrics (n. of cases per case of the prior rank), the numbers observed for different phyla reveal very variable individual trends. Fig. 12 reports a number of selected examples in which the same parameter that has been described for the aggregate taxa within a domain, or by pooling lineages under a species abundance criterion in the previous graph, is instead applied to the numbers that regard the situations of single phyla The cases shown in Fig. 12 reveal a set of rather different patterns. These span from the ‘orders bloom’ of the Pseudomonadota, to the ‘classes-first’ outcome of the Actynomicetota and Planctomycetota or to the variety within Gram+ with the ‘peaking at orders’ for Bacillota (the phylum that includes *Lactiplanctibacillus*), or with the added ‘peaking at genera’ hump for Bacillota A (the phylum that includes *Clostridium*), or for the peaking at family rank for the pathogen Campylobacterota.

**Fig. 12.**
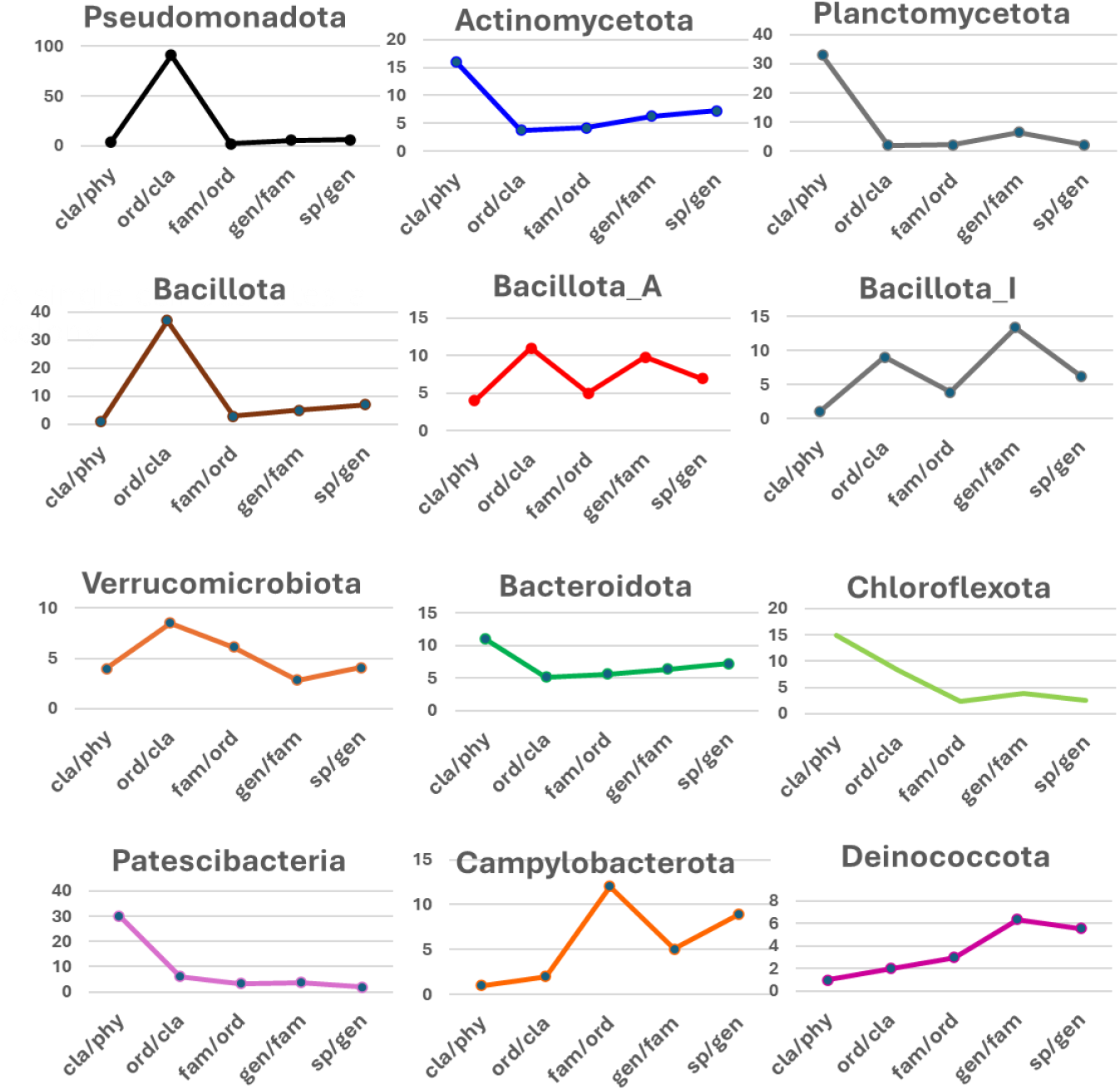
Evolutionary efficiency rates at the phylum level. Twelve phyla were chosen as representative examples of different outcomes in building the ranks content.

### A closer look (species-level) at some popular microbes

The graphs assembled in Fig. 13 report in the histogram bars the numbers of classes, orders, families, and genera in the lineage of each chosen case, as well as the number of species shared within its genus. Notice the Order-& genera-rich case of *L. plantarum*, which deviates from the global style of its own phylum Bacillota, that showed just the order peak (Fig. 12). A similar trend is seen for *E*.*coli. S. griseus* is instead within the most species-rich genus of the whole bacterial taxonomy (*Streptocomyces*, with its 1,070 species). The cheese starter *Streptococcus thermophilus* is within another particularly species-endowed genus case. Notice also the class- and family-rich but species-poor extremophile *Sulfolobus*, and, in pathogens, the radiative taxonomy gradient of *H. pylori*, which is in line with the rise of its animal hosts. In fact, being animals recent in natural history, little deep branching (few classes) and more recent speciation would be expected for an animal pathogen as *H. pylori* current taxonomic tree.

**Fig. 13.**
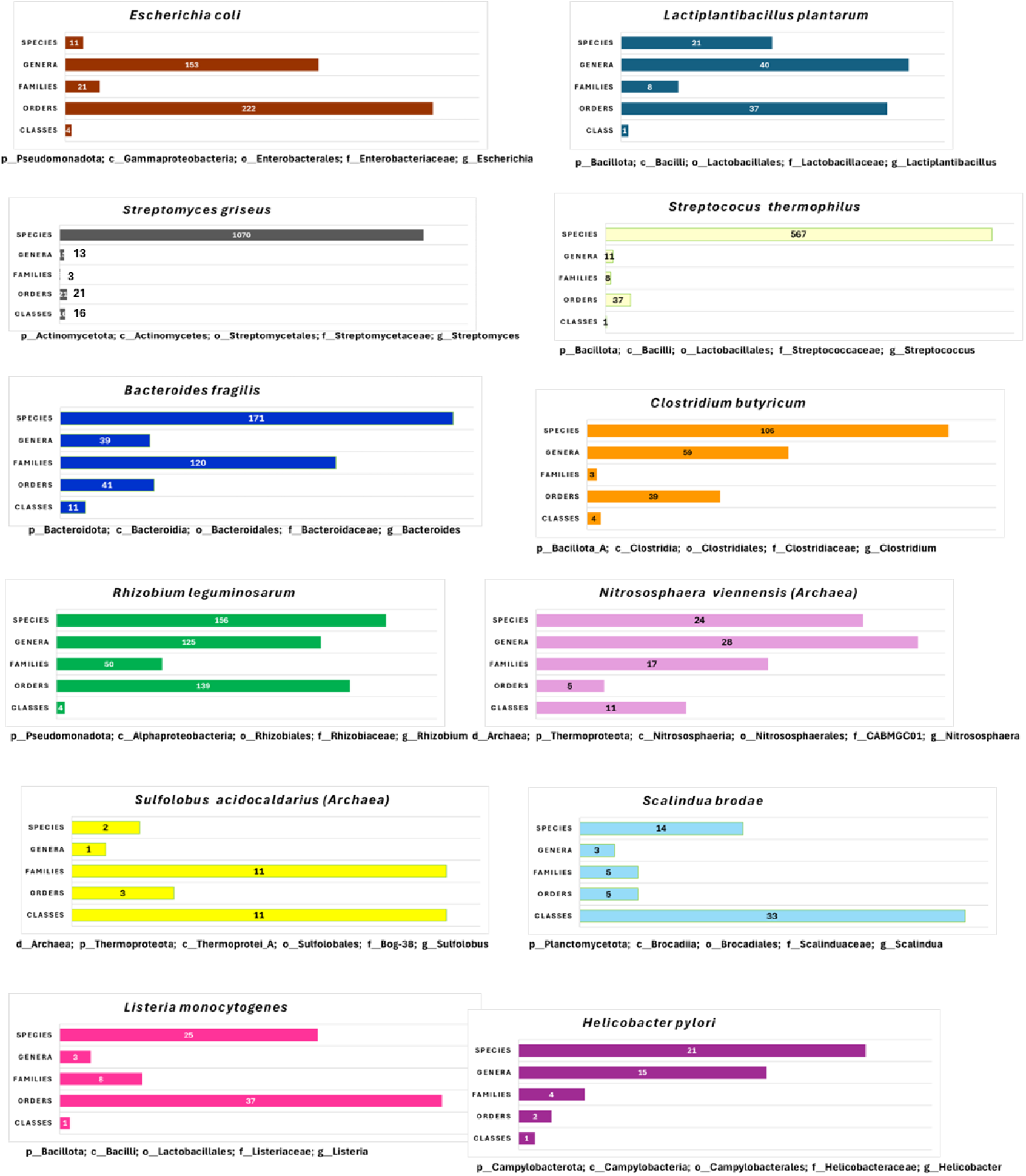
Numbers of members of each rank for the lineages of 12 characteristic examples. The horizontal histogram bars show the numbers of classes, orders, families, and genera featured by each chosen lineage, and the number of other species with which the one of each example shares its presence within the genus.

### Conclusive remarks

A general reminder is needed before reflecting on any data of this kind. When looking back towards the origin of a species within one of the two prokaryotic domains and walking along its lineage from finer to coarser ranks, one might lean, deceptively, towards the belief that phyla should be the oldest taxonomic ‘boxes’, while species would be the newest. This concept can be handy when describing the directional process from the ranks’ hierarchy point of view, and, for convenience, it has been applied in some points, also when commenting on the results of the present analysis. However, the situation should actually be seen as the opposite. Phyla are classically defined by a fork of divergence among each other in a reference gene, which has <u>today</u> mutated enough to reach the cutoff levels that we envisage as boundaries for the largest ranks of a domain. But this is indeed the achievement that those groups have earned <u>to date</u>, after all the erosion of similarity that has been brought by the passage of time. Vice versa, when the two ancestors of two bacteria that, at present, are classified into two different phyla, started to diverge from the latest common ancestor of both, they did not embody two phyla and yet not even two species as, initially, their difference in the reference gene would have been a single nucleotide, and in an about 1500 bases-long 16S rRNA it takes at least 38 nucleotides to reach the less-than-97.5% homology that sets the species border. A single mutation in the 16S rRNA gene has been estimated to occur every 4.4 million years, which sets to about 166 million years the time to achieve enough dissimilarity to split apart two species (Clark et al., 1999). Therefore, in bacteria that are now ascribed to different phyla, the ‘old fact’ that occurred is not the birth of their phyla but just the branching, which, deep in the past, started a variation in one of their sequences. Phyla are therefore the youngest ranks that have finally mathematically arisen, having gained enough differences to exit from a common class-level container. Being life traced back to a single origin, therefore, also by looking at natural history in the time-forward mode, that single ancestor could not belong to any categorizable phylum nor to any rank at all, as our metrics in systematics define each of those sets (the ranks) by the extent of lost sequence homology. All our ranks concepts are thus defined *a posteriori* and, conceptually, only after millions of years since a phylogenetic branch started, one could have slowly witnessed the ‘existence’ of species, then that of genera, then families, orders, and so on, to accumulate enough percentage of differences to finally support the grounds to define classes and then, at last, phyla. This also implied that further deeper and higher ranks beyond phyla and domains could be coined in a future in which a higher divergence could accommodate their existence. That future has actually already begun, as, in fact, in Prokaryotes, between domain and phylum, the Kingdom rank has been recently placed, and four bacterial kingdoms (Bacillati, Fusobacteriati, Pseudomonadati, Thermotogati), and three archaeal kingdoms (Methanobacteriati, Nanobdellati, Thermoproteati) have been designated, including also, for some of those, additional extra ranks like the Subkingdom and the Superphylum (Göker and Oren, 2024).

But, no matter how many cutoff points we will introduce, in practice, all the taxonomy architecture could be seen as a virtual hierarchical net of terms (that are cast over something that is actually a continuum). Undeniably, it is an extremely useful framing cabinet to distinguish the microbial life forms, but, as regards evolution, it is a system that uses name-containing sets that can be just coined backwards, in retrospect, ‘after the race’.

The presentation of the results of this analysis and of the comments stemming from their observation are meant to give a brief and non-exhaustive series of examples on how the data hereby provided as deliverable (supplementary material dataset S1) can be used in multiple ways and enable the study of individual details from any of the 107,235 species featured.

Moreover, the updates in the bacterial and archaeal genomes database at GTDB will allow the expansion and refinement of the information that can be extracted from their further releases with the same protocol described in the method section of this article.

In essence, the mathematical elaborations done in the present data analysis show that several clues about microbial evolution are concealed in plain and available datasets and can be mined and interpreted using very simple methods.

## Supporting information

Supplementary Dataset S1 Bacterial and Archaeal elaborated counts

